# Small RNAs are trafficked from the epididymis to developing mammalian sperm

**DOI:** 10.1101/194522

**Authors:** Upasna Sharma, Fengyun Sun, Brian Reichholf, Veronika A. Herzog, Stefan L. Ameres, Oliver J. Rando

## Abstract

RNAs present in mature mammalian sperm are delivered to the zygote at fertilization, where they have the potential to affect early development. The biogenesis of the small RNA payload of mature sperm is therefore of great interest, as it may be a target of signaling pathways linking paternal conditions to offspring phenotype. Recent studies have suggested the surprising hypothesis that the small RNA payload carried by mature sperm may include RNAs that were not synthesized during testicular spermatogenesis, but that are instead delivered to sperm during the process of post-testicular maturation in the epididymis. To further test this hypothesis, we characterized small RNA dynamics during testicular and post-testicular germ cell maturation in mice. We show that purified testicular germ cell populations, including mature testicular spermatozoa, carry extremely low levels of tRNA fragments (tRFs), and that tRFs become highly abundant only after sperm have entered the epididiymis. The process of small RNA delivery to sperm can be recapitulated in vitro, as caput epididymosomes deliver small RNAs including tRFs and microRNAs to mature testicular spermatozoa. Finally, to definitively identify the tissue of origin for small RNAs in sperm, we carried out tissue-specific metabolic labeling of RNAs in intact mice, finding that mature sperm carry small RNAs that were originally synthesized in the somatic cells of the epididymis. Taken together, our data demonstrates that soma-germline small RNA transfer occurs in male mammals, most likely via vesicular transport from the epididymis to maturing sperm.

## INTRODUCTION

Inheritance of phenotypic changes in the absence of any change in DNA is known as epigenetic inheritance, and is essential for cell state inheritance during mitosis. Epigenetic information is not only inherited during cell division, but at least some epigenetic information can be transmitted from one generation to the next in multicellular organisms, a mechanistically complex process that requires epigenetic information to be maintained throughout the disruptive processes of gametogenesis, fertilization, and early embryonic development. Although decades of study of cell state inheritance and other cases of epigenetic inheritance have identified several major molecular conduits of epigenetic information – chromatin structure, small RNAs, DNA modifications, prions, and transcription factors – much remains to be understood regarding the establishment and erasure of epigenetic information during gametogenesis and early development.

Small (<40 nt) RNAs are one of the best-established carriers of epigenetic information, and play key roles in epigenetic inheritance paradigms in *C. elegans* (Fire et al., 1998), *A. thaliana* (Arteaga-Vazquez and Chandler, 2010; Hamilton and Baulcombe, 1999; Zilberman et al., 2003), *S. pombe* (Volpe et al., 2002), *D. melanogaster* (Brennecke et al., 2008; de Vanssay et al., 2012), and mammals (Chen et al., 2016; Gapp et al., 2014; Rassoulzadegan et al., 2006; Rodgers et al., 2015; Sharma et al., 2016). Multiple distinct families of small RNAs have been described which differ from one another in their biogenesis and in their regulatory functions, and include well-studied species such as microRNAs, endo-siRNAs, and the gamete-enriched class of piRNAs which are crucial for self-nonself recognition in the germline (Aravin et al., 2006; Batista et al., 2008; Brennecke et al., 2007; Brennecke et al., 2008; Mochizuki et al., 2002; Shirayama et al., 2012; Vagin et al., 2006). In addition, a number of less-studied small RNA species have been described, including small RNAs derived from various longer noncoding RNAs including tRNAs – these latter RNAs are most commonly referred to as tRNA fragments, or “tRFs” (Anderson and Ivanov, 2014; Sobala and Hutvagner, 2011). While tRFs have been found in numerous small RNA sequencing datasets, they appear to be particularly abundant in contexts related to reproduction, as for example 3’ tRFs are the primary cargo of the *Tetrahymena* Piwi-related argonaute Tiwi12 (Couvillion et al., 2010), while 5’ tRFs comprise the vast majority (~80%) of small RNAs in mature mammalian sperm (Chen et al., 2016; Garcia-Lopez et al., 2014; Peng et al., 2012; Sharma et al., 2016).

The mechanisms responsible for the biogenesis of the small RNA payload of mammalian sperm are not completely understood. Small RNA dynamics have been extensively studied during the process of testicular spermatogenesis, where three piwi-clade argonaute proteins (MIWI, MILI, and MIWI2) orchestrate two major waves of piRNA biosynthesis (Aravin et al., 2006; Aravin et al., 2007; Kuramochi-Miyagawa et al., 2004; Li et al., 2013). Initially, “pre-pachytene” piRNAs – comprised primarily of piRNAs derived from repeat elements – are expressed beginning in spermatogonia (Aravin et al., 2007), followed by a wave of unique pachytene piRNAs which are processed from 214 long noncoding precursor transcripts (Li et al., 2013). Given that small RNAs found in testes are almost entirely comprised of piRNAs, it is curious that the vast majority of small RNAs present in mature sperm are 5’ tRNA fragments, as noted above. Indeed, 5’ tRFs are vanishingly scarce (~2% of all small RNAs) in intact testes despite the fact that testes are primarily comprised of gametes at varying stages of development, raising the broader question of how and when small RNAs are gained or lost during post-testicular maturation.

We recently discovered a potential role for the epididymis – a long convoluted tubular structure where mammalian sperm undergo post-testicular maturation – in modulating the small RNA repertoire of maturing sperm (Sharma et al., 2016). We found that tRNA cleavage occurs robustly in the epididymis, and that small vesicles secreted by the epididymal epithelium, known as epididymosomes (Belleannee et al., 2013; Krapf et al., 2012; Sullivan et al., 2007; Sullivan and Saez, 2013), carry a very similar population of small RNAs to that gained by sperm during epididymal transit (Sharma et al., 2016). Moreover, as sperm successively pass from caput (proximal) to cauda (distal) epididymis, sperm gain those specific small RNAs that are locally enriched in the underlying epithelium. Finally, we and others (Reilly et al., 2016) have shown that epididymosomes can deliver small RNAs to relatively immature caput epididymal sperm in vitro, suggesting that the sperm small RNA repertoire is shaped by soma-to-germline trafficking of small RNAs.

Here, we rigorously test this hypothesis, characterizing developmental dynamics of small RNAs across multiple purified sperm populations spanning testicular and post-testicular development. We detected very low levels of tRFs in all testicular germ cell populations, including mature testicular spermatozoa, confirming that tRFs are only gained upon entry into the epididymis. This maturation step could be partially recapitulated in vitro, as we show that epididymosomes from the proximal epididymis can deliver a wide variety of small RNAs, including both microRNAs and tRFs, to testicular spermatozoa. Finally, we use a chemogenetic approach to definitively identify the tissue of origin for small RNAs present in mature cauda epididymal sperm. Using epididymis-specific expression of UPRT to enable 4-thiouracil labeling of small RNAs synthesized in the caput epididymis, we show that mature cauda sperm carry small RNAs first synthesized in the epididymis epithelium. Taken together, these data conclusively demonstrate that small RNAs are trafficked from soma to germline in mammals, and provide further evidence that the small RNA payload of mature sperm is shaped by successive waves of epididymosomal delivery during epididymal maturation.

## RESULTS

### Small RNA dynamics during sperm development and maturation

We and others have previously reported that sperm isolated from either the cauda (distal) or caput (proximal) epididymis carry a small RNA payload comprised primarily of 5’ tRNA fragments (tRFs). In contrast, tRNA cleavage as assayed both by small RNA sequencing and by Northern blot is nearly undetectable in intact testes, where the vast majority of small RNAs are piRNAs (Peng et al., 2012; Sharma et al., 2016). Although these data suggest that tRFs are gained upon entry into epididymis, it remains formally possible that tRFs are gained during the last stages of testicular spermiogenesis as round spermatids mature into testicular spermatozoa.

To better characterize small RNA dynamics throughout the processes of testicular spermatogenesis/spermiogenesis and post-testicular maturation, we purified six germ cell populations from the reproductive tracts of sexually mature (8-12 weeks) male mice. We purified four testicular populations, including primary spermatocytes and two fractions of round spermatids as in (Sharma et al., 2016), as well as testicular spermatozoa which were purified using percoll gradients (**Materials and Methods, Supplemental Figure S1**). In addition, mature sperm were obtained from caput and cauda epididymis as previously described. Total RNAs were extracted and small RNAs (18-40 nucleotides) were purified and subject to deep sequencing. Our data recapitulate prior findings, with piRNAs dominating the small RNA repertoire of spermatocytes and round spermatids, and tRFs being the primary small RNA population in both caput and cauda epididymal sperm (**Figure 1, Table S1**). Turning to the previously-unexamined testicular spermatozoa, we find that even at this late stage of maturation sperm resemble other testicular populations rather than epididymal sperm – they continue to carry high levels of piRNAs (~70% of small RNA reads), having yet to gain abundant tRFs (~7% of reads). These data provide further insight into a major transition in sperm small RNAs, demonstrating that the RNA payload of mammalian sperm shifts from piRNAs to tRFs at some point after exiting the testis for the epididymis.

**Figure 1:**
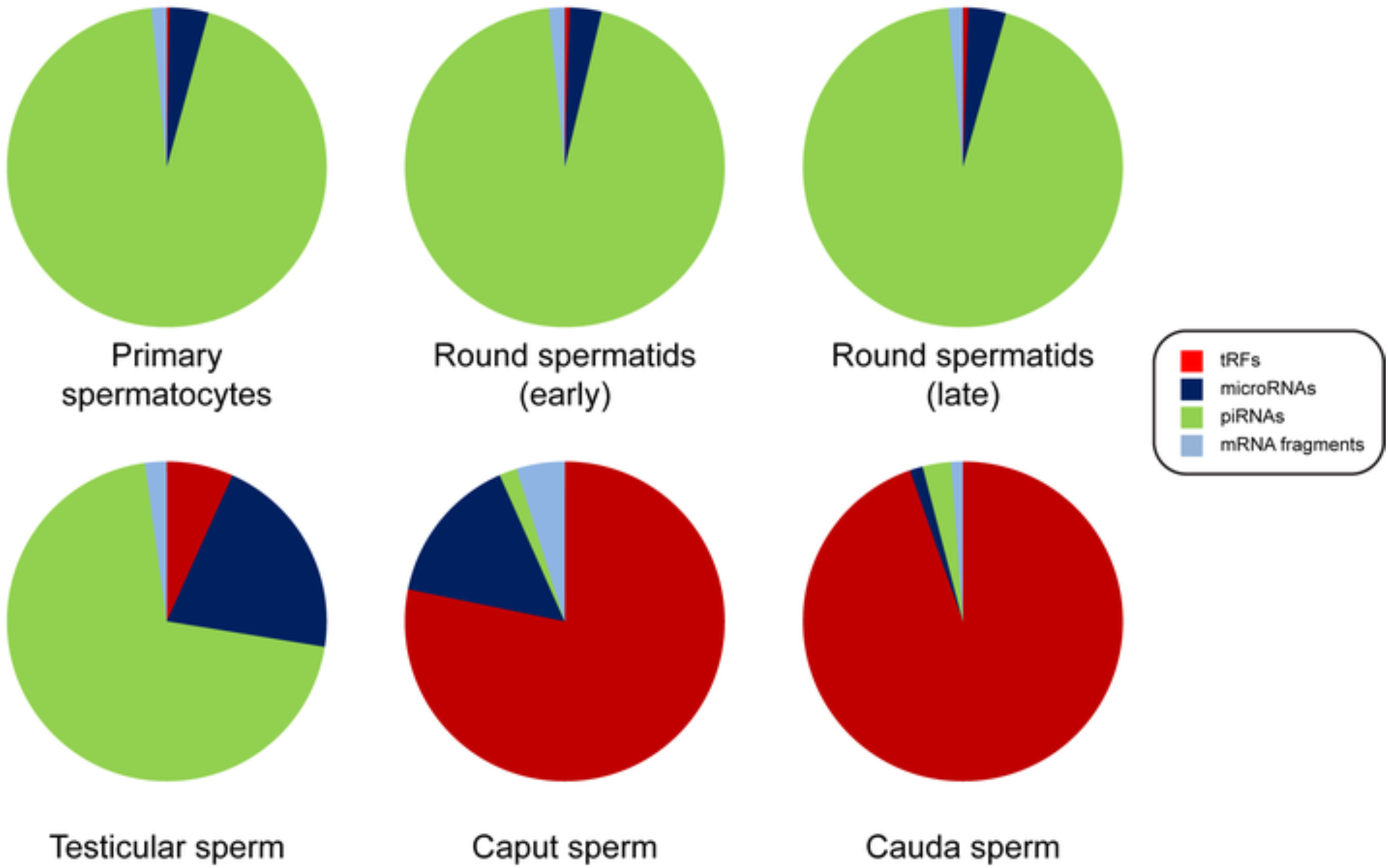
Bulk changes to small RNA payload throughout testicular and post-testicular sperm development. Small RNAs (18-40 nt) were isolated from the indicated germ cell populations and subject to deep sequencing. Deep sequencing reads were mapped to rRNAs, tRNAs, miRbase, Repeatmasker, to unique piRNA precursors, and to RefSeq. After removing reads mapping to rRNAs, small RNA abundance was normalized to parts per million of remaining mapped reads, and pie charts show fraction of small RNAs mapping to the indicated small RNA categories(reads mapping to Repeatmasker and to unique piRNA precursors are merged into the piRNA category).

In addition to the bulk differences in abundance of broad classes of small RNAs, we also find marked differences in specific individual RNA species present in each germ cell population, confirming for example the previously-described gain in tRF-Val-CAC that occurs as sperm transit from caput to cauda epididymis (Sharma et al., 2016). In order to broadly illuminate the major dynamic behaviors observed during sperm maturation, we grouped small RNAs across the six stages studied here using K-means clustering (**Figure 2A,** K=6). This visualization highlights the major transitions in small RNA populations described above, as Clusters 1 and 2 show the global loss of piRNAs that occurs after sperm exit the testis, and Cluster 6 illustrates the gain of tRFs that occurs in the epididymis. Interestingly, a handful of other small RNAs exhibit similar behaviors to piRNAs and tRFs, as for example miR-34c is highly expressed in testis but is largely jettisoned upon entry into the epididymis. Other clusters reveal unanticipated behaviors, as for example Cluster 3 – characterized by maximal expression in testicular sperm – includes a relatively large number of piRNAs (all of which represent the unique “pachytene” piRNA class, rather than deriving from repeat elements), thus identifying a wave of piRNA biogenesis that occurs during the very last stages of testicular sperm development. Cluster 5 is also notable, as it is primarily comprised of fragments of mRNAs that are highly-expressed in the caput epididymis (Johnston et al., 2005).

**Figure 2:**
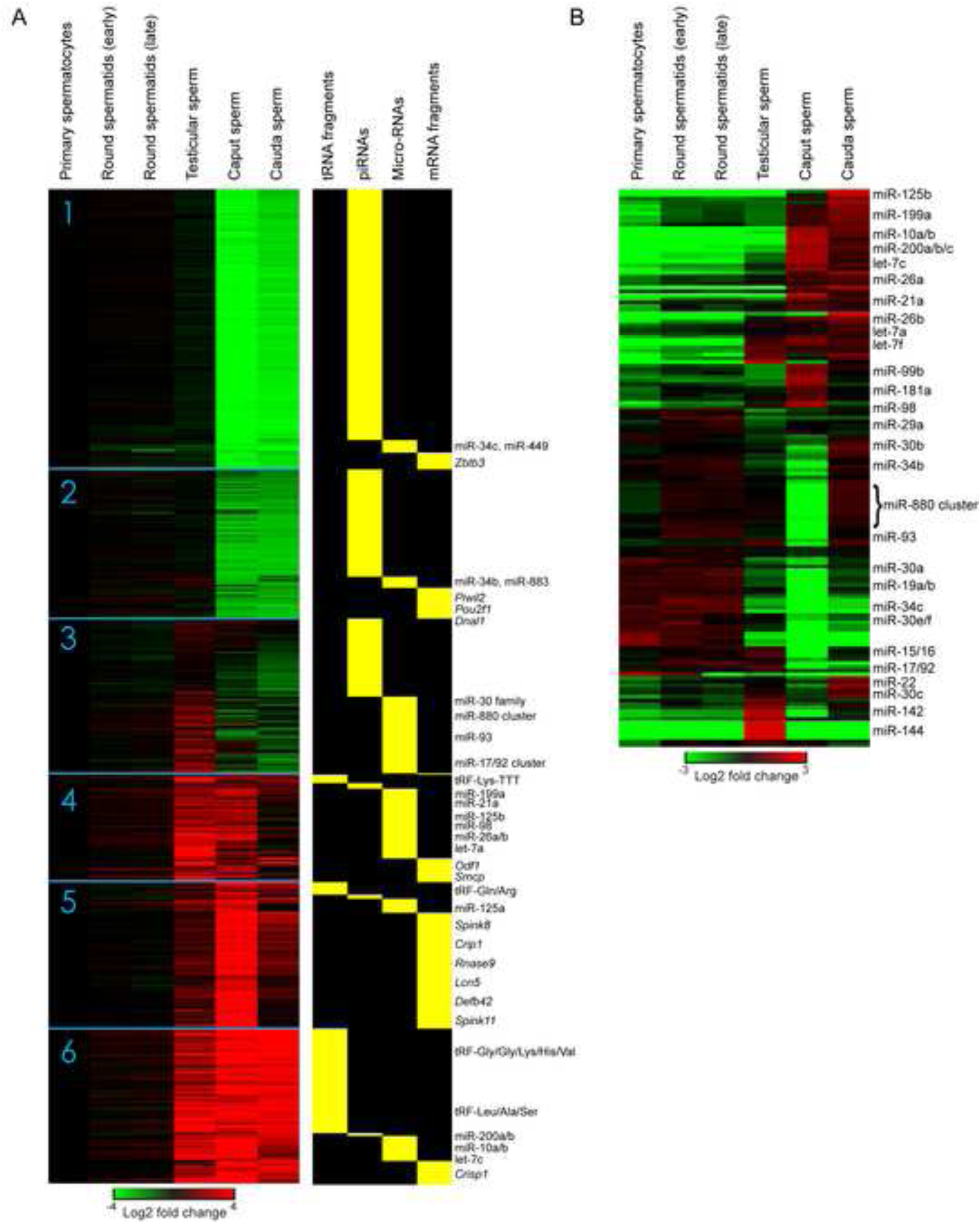
Detailed small RNA dynamics across sperm development. Comparative analysis of small RNAs in the six developmental stages of sperm development depicted in the form of heat maps. A) K means cluster (K=6) showing all small RNAs with a maximum abundance of at least 50 ppm at one developmental stage. Data are normalized first to ppm (all non-rRNA mapped reads) for a given developmental stage, then normalized to their abundance in primary spermatocytes. Left panels show change in abundance relative to spermatocytes, right panels show small RNA annotations. B) MicroRNA dynamics during sperm development. Here, microRNA abundances were re-normalized only to the total microRNAs at a given developmental stage, then zero-centered – re-normalized according to the average abundance in the six stages analyzed.

In contrast to the relatively tissue-specific expression of piRNAs and tRNA fragments, microRNAs exhibit a more diverse range of behaviors during spermatogenesis and sperm maturation. However, the visualization in **Figure 2A** is dominated by the bulk changes in piRNAs and tRFs, complicating the interpretation of microRNA dynamics – for example, the apparent restriction of Cluster 3 microRNA expression to testicular sperm might simply reflect diminished small RNA complexity in this sperm population, which has started to lose piRNAs but has yet to substantially gain tRFs (**Figure 1**).

To investigate microRNA dynamics in more detail, we renormalized miR abundances according to total miR levels for each sperm population, rather than to all small RNAs, to avoid the confounding normalization effects of bulk piRNA/tRF loss and gain. This analysis revealed a range of dynamic behaviors of individual microRNAs (**Figure 2B**). The majority of microRNAs are either expressed during testicular spermatogenesis and then lost/diluted at later developmental stages, or conversely are expressed/gained primarily in later stages of sperm maturation. As example of the former behavior, microRNAs such as miR-30a and miR-34c are lost during epididymal maturation, presumably due to degradation or elimination in the residual body shed by sperm in the epididymis (Sprando and Russell, 1987). A subset of microRNAs such as miR-144 and miR-142 are highly abundant only in testicular spermatozoa, and are presumably only expressed during late stages of spermatogenesis and then lost by a similar mechanism. The converse behavior, in which miRNAs such as miR-125b, miR-199a, miR-200b/c, miR-10a/b and let-7 family members increase in relative abundance during epididymal maturation, is consistent with these RNAs potentially being trafficked to epididymal sperm alongside the bulk delivery of tRFs (Reilly et al., 2016; Sharma et al., 2016).

More surprising are cases of microRNAs that are abundant in testicular sperm as well as cauda sperm, but which appear to be scarce specifically in caput sperm. Interestingly, the majority of such microRNAs originate from genomic microRNA clusters of various sizes, ranging from small clusters (miR-15/16, the “oncomiR”17-92 cluster, etc.) up to the megabase-scale imprinted X-linked miR-880 cluster. This behavior is most consistent with these microRNAs initially being lost (by degradation or elimination) upon epididymal entry, followed by a second load of these miRNAs either being generated *in situ* in cauda sperm from their precursor molecules, or shipped from epididymis epithelium to the mature sperm by epididymosomes. Consistent with the latter hypothesis, many of the microRNAs that are specifically scarce in caput sperm are highly abundant in cauda epididymosomes (**Table S1**). These data thus uncover surprisingly complex small RNA dynamics during sperm development, and further underscore the major role of epididymal transit in modulation of the sperm epigenome.

### Caput epididymosomes deliver small RNAs to testicular spermatozoa

Our data confirm and extend prior reports showing major differences between the small RNAs generated during testicular spermatogenesis and those gained during epididymal transit. What is the mechanism by which tRFs and other small RNAs are gained as sperm enter the epididymis? We previously showed that epididymosomes secreted by the epithelium of cauda epididymis have a similar RNA payload to that of mature sperm, and we and others showed that epididymosomes can deliver small RNAs to relatively “immature” caput sperm (Reilly et al., 2016; Sharma et al., 2016). However, as the RNA repertoire of caput sperm is not particularly distinct from that of cauda sperm – caput sperm already carry high levels of most tRNA fragments, for example – these reconstitution studies could only focus on a small handful of fairly cauda-specific species such as tRF-Val-CAC. In contrast, the RNA payload of testicular spermatozoa, which is dominated by piRNAs, is highly distinct from that of either epididymal sperm population, providing us with a “blank slate” to more broadly explore the ability of epididymosomes to deliver RNAs to immature sperm.

Testicular sperm were isolated, then were either mock treated or were incubated for two hours with caput epididymosomes, and resulting sperm were extensively washed to remove any contaminating vesicles (**Figure 3A**). We first used TaqMan qRT-PCR assays to ask whether two prominent tRFs present in epididymosomes but scarce in testicular sperm could be delivered by this reconstitution, finding that epididymosomal delivery results in a ~4-10-fold increase in abundance for these two tRFs (**Figure 3B**). To more broadly interrogate epididymosomal RNA delivery to sperm, we next carried our deep sequencing of small RNAs from mock and “reconstituted” sperm. Globally, we observed significantly higher levels of tRFs and microRNAs in reconstituted spermatozoa, relative to mock-treated sperm (**Figures 3C, Table S2**). Although two hours incubation is clearly insufficient to fully establish the caput small RNA payload, this is perhaps unsurprising given that purified caput sperm from intact animals have spent several days undergoing this delivery process.

**Figure 3:**
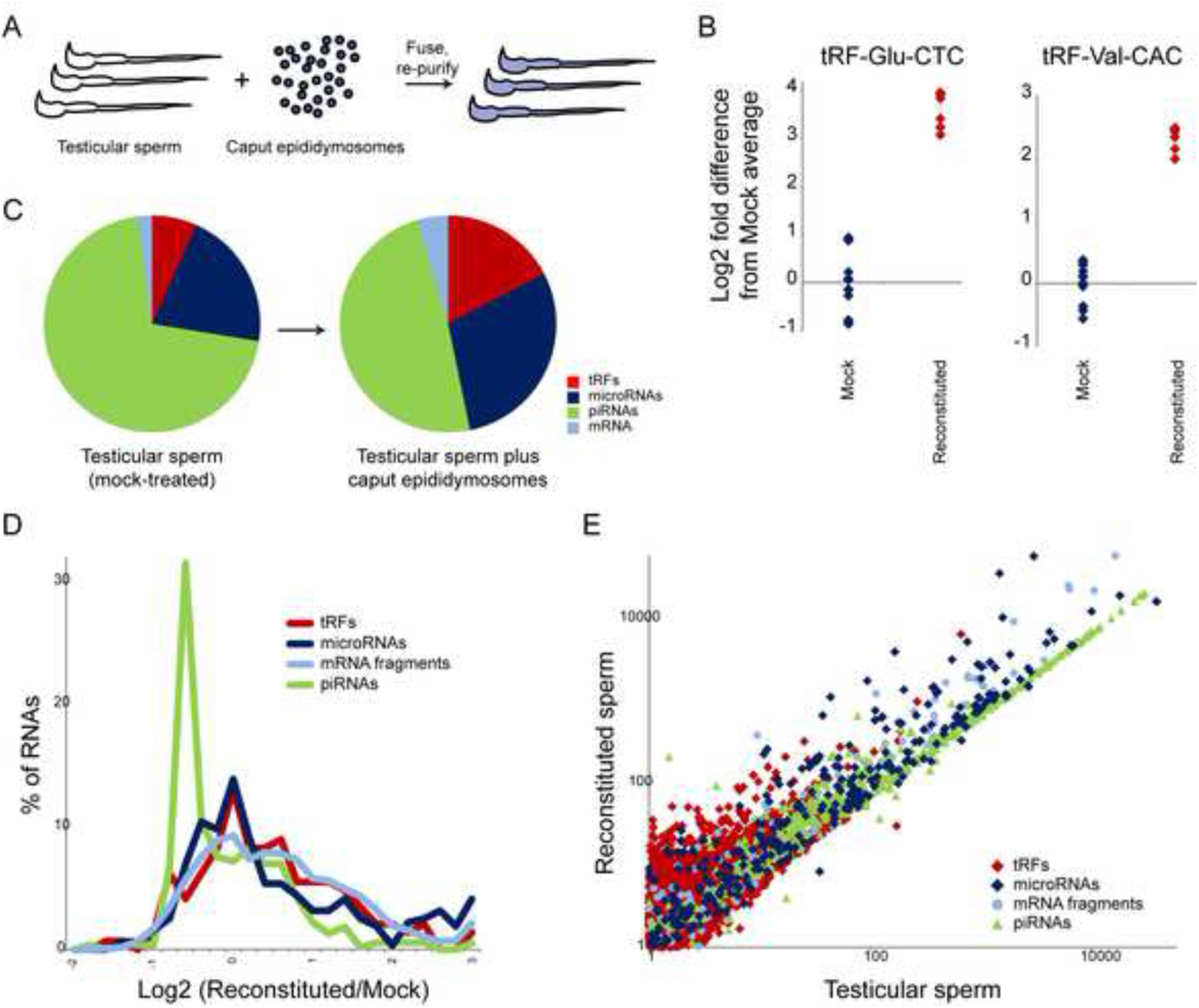
Reconstitution of small RNA delivery to testicular sperm. A) Experimental schematic. Purified testicular sperm, which carry extremely low levels of tRFs, were incubated with caput epididymosomes for 2 hours, followed by extensive washing. Small RNAs were purified from either mock-treated testicular sperm, or reconstituted sperm, and deep sequenced. B) Delivery of two prominent tRFs to testicular sperm. Taqman q-RT-PCR for the indicated tRFs, with individual replicates plotted (on a log2 y axis) relative to the average level present in Mock-treated testicular sperm. C) Deep sequencing of testicular sperm either mock-treated or incubated with caput epididymosomes. Pie charts show average levels of various small RNA classes, revealing increased levels of microRNAs and tRFs delivered to testicular sperm by epididymosomes. D) Distribution of small RNA changes in sperm following epididymosome fusion. X axis shows the log2 fold difference between reconstituted and mock-treated sperm – positive values indicate delivery by epididymosomes. Importantly, because genome-wide normalization methods assume no global change in RNA abundance, RNAs that are in fact unaltered during a global gain of RNAs will exhibit a decrease in *relative* abundance, as observed here for the bulk of piRNAs (average log2 fold decrease of~0.6). Values over -0.6 therefore indicate gain of RNA during the fusion protocol. E) Scatterplot of small RNA abundances from deep sequencing data. The strong diagonal for piRNAs again indicates RNAs present in testicular sperm that are not affected by the delivery process. Note that essentially all RNAs here either lie along this diagonal, indicating that they are either absent or nearly so in caput eididymosomes, or above the diagonal, indicating widespread delivery of many RNA species to testicular sperm during reconstitution.

More granular analysis of specific small RNA species revealed that the vast majority of piRNAs were unaffected by epididymosomal fusion (**Figures 3D-E**), consistent with the fact that piRNAs are expressed during testicular spermatogenesis and are largely absent from epididymosomes (and indicating that epididymosomal fusion does not induce piRNA degradation). This is clearly visualized in the scatterplot in **Figure 3E**, where piRNA abundance is strongly correlated between mock-treated and reconstituted sperm samples. Globally, nearly all small RNAs either fall on the same diagonal as the piRNAs – meaning that they are unaffected by epididymosomal fusion – or above this diagonal, consistent with delivery by epididymosomes. Individual tRFs and microRNAs exhibited a range of behaviors (**Figures 3D-E**), ranging from those unaffected by epididymosomal incubation and which are generally already abundant in testicular sperm and/or absent from caput epididymosomes, to efficiently-delivered small RNAs that are highly abundant in caput epididymosomes (such as miR-10a/b, miR-148, miR-143, tRF-Glu-CTC, tRF-Gly-GCC, and tRF-His-GTG – **Table S2**).

Overall, these data show that caput epididymosomes are capable of fusing with mature testicular spermatozoa to deliver their small RNA cargo in vitro, and together with the in vivo small RNA dynamics detailed above are most consistent with a mechanism of RNA biogenesis in mammalian sperm in which small RNAs generated in the epididymis are trafficked to sperm in epididymosomes.

### Chemogenetic tracking of small RNAs from epididymis to sperm

Although multiple observations here and elsewhere are most readily explained by the hypothesis that small RNAs in mature sperm are first generated in the epididymal epithelium, we have not formally ruled out the possibility that the small RNAs gained by sperm during epididymal maturation in intact animals are instead processed *in situ* from precursor molecules first synthesized during testicular spermatogenesis. In order to distinguish between these possibilities, we developed a strategy to label and track small RNAs synthesized in specific cell types. We made use of the “TU tracer” mouse (Gay et al., 2013), in which tissue-specific expression of the Cre recombinase is used to drive expression of the gene encoding uracil phosphoribosyltransferase (UPRT) (**Figure 4A**). Expression of UPRT allows cells to incorporate 4-thiouracil (4-TU) into newly-synthesized RNAs, thereby enabling tissue-specific metabolic labelling of RNA (**Figure 4B**).

**Figure 4:**
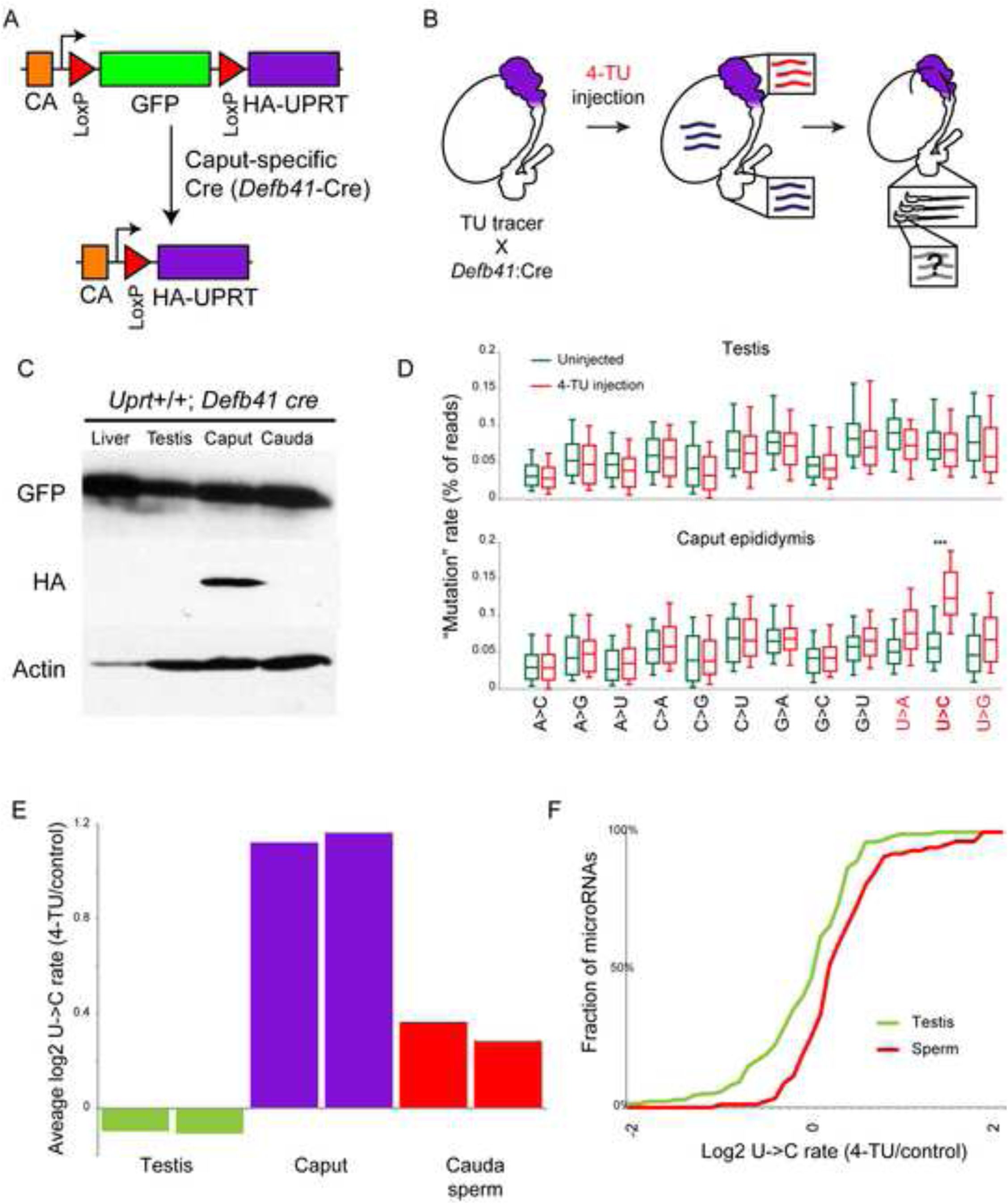
Chemogenetic tracking of RNAs from epididymis to sperm. A) Schematic of the TU tracer locus. In the absence of Cre, GFP is expressed from a ubiquitous promoter, with no UPRT expression. Cre drives LoxP recombination, eliminating GFP and juxtaposing the promoter with HA-UPRT. B) Experimental design. UPRT is expressed specifically in the initial segment of the caput epididymis due to expression of *Defb41*:Cre. Upon injection of 4-thiouracil, RNAs synthesized in this tissue incorporate 4-TU. Animals are injected with 4-TU every 2 days for 9 days (5 injections), and 5 hours after injection on Day 9 sperm are isolated from the cauda epididymis and assayed for 4-TU labeling of small RNAs. C) Western blots showing HA-UPRT expression specifically in the caput epididymis of *Defb41*:Cre X TU-tracer animals. D) SLAM-Seq induces miscloning of uracil in a Cre and 4-TU-dependent manner. *Defb41*:Cre X TU-tracer animals were either injected with 4-TU (every other day for 9 days) or sham-injected, and testis and caput epididymis were dissected. Small RNAs were purified and subject to SLAM-Seq. Reads were mapped to microRNAs using a mismatch-tolerant custom pipeline (**Methods**) and nucleotide misreads were compiled for all microRNAs with at least 1000 reads (boxes show median and 25^th^/75^th^ percentiles, whiskers show 5^th^ and 95^th^ percentiles). 4-TU induces misreading of uracils specifically in Cre-expressing tissue, with U->C being the most common mutation observed (p = 1.2e-6, t test). E) Log2 ratio of U->C rates for 4-TU injected vs. uninjected animals, for all microRNAs with at least 1000 reads for the indicated tissues. Two bars show two biological replicate experiments. U->C mutation rate increases in caput epididymis, as expected (see panel D), and also in cauda sperm, indicating the presence of caput-derived microRNAs in cauda sperm. The modest increase in U->C mutation in mature sperm reflects multiple factors: 1) the signal to noise in caput epididymis is already modest (2-fold); 2) cauda sperm presumably carry a wide range of unlabelled RNAs originating in the testis or other parts of the epididymis; 3) sperm in the cauda epididymis can survive for longer than the time frame of this experiment and so will include a population of unlabelled sperm that passed through the caput epididymis prior to 4-TU injections. F) Cumulative distribution plots for all individual microRNAs (>1000 reads) in the testis and cauda sperm dataset, as indicated. The right shift in the curve for sperm indicates significant (KS p = 0.0003) enrichment of 4-TU-induced mutations, relative to the testis control. See also **Supplemental Figure S4B**.

To assess the efficiency and specificity of this system, we first generated mice expressing UPRT specifically in liver by mating *Albumin*-Cre (*Alb*-Cre) mice with TU-tracer mice. Cre-mediated recombination was extremely efficient, with nearly quantitative deletion of the GFP insert and concomitant expression of UPRT (**Supplemental Figure S2A**) – remaining GFP likely reflects non-hepatocyte cell populations, such as blood cells, present in the liver. The resultant mice expressed UPRT specifically in liver (**Supplemental Figure S2A**), and injection of 4-TU in these mice resulted in liver-specific labeling of RNAs, with undetectable background incorporation in other tissues (**Supplemental Figures S2B-C**). Northern blot of labeled RNAs revealed 4-TU labeling of a wide range of RNA species including microRNAs (see below), but tRNAs and rRNAs were not efficiently labeled (not shown), either due to poor use of 4-TU by PolI and PolIII, or due to checkpointing of these typically highly-modified RNAs. This protocol therefore enables us to characterize microRNA trafficking in vivo, but unfortunately cannot be used to track tRFs.

Although tracking TU-labeled RNAs in vivo could in principle be accomplished by biotinylation of 4-TU (**Supplemental Figures S2B-C**) and deep sequencing of avidin-purified RNAs, we note that even in an abundant tissue such as the liver a very small fraction of RNA was 4-TU-labeled, and the abundance of any RNAs trafficked between cells would of course be far lower. Along with the fact that sperm overall have very low levels of total RNA, we anticipated extreme difficulty in purification of 4-TU-labeled RNAs. We therefore turned instead to thiol(S)-linked alkylation for the metabolic sequencing of RNA (SLAM-Seq, Herzog *et al*., submitted – **see accompanying manuscript**) for characterization of 4-TU-labeled small RNAs. This method is based on chemically modifying the 4-thiouracil in RNA with iodoacetamide to generate a covalent adduct that causes nucleotide misincorporation during reverse transcription, with G misincorporation opposite the modified U (seen as a U->C mutation in downstream deep sequencing) occurring most frequently. This method enables quantitative analysis of 4-TU incorporation in any clonable population of small RNAs, avoiding potentially lossy purification. To quantitate the signal-to-noise for this method, we first applied SLAM-Seq to small RNAs purified from the liver of *Alb*:Cre-expressing mice (**Supplemental Figure S3**). Background mutation rates in the cloning process were characterized using *Alb*:Cre mice that had not been injected with 4-TU, while analysis of cauda epididymis in these animals controlled for both cloning errors and for 4-TU incorporation into non-UPRT-expressing cells. As expected, we find that 4-TU incorporation caused a significant increase in misincorporation events specifically opposite U, with the strongest effect on U->C transitions (**Supplemental Figure S3**). This required 4-TU injection, was dose-dependent, and required tissue-specific expression of UPRT. Variation in the extent of U->C misincorporation for individual microRNAs reflects in vivo RNA synthesis and decay rates, as well as any effects of local sequence context on RT misincorporation. Together, these results demonstrate our ability to identify RNAs first synthesized in a given genetically-defined cell type.

We therefore applied this system to track microRNAs synthesized in the caput epididymis, using the *Defb41* promoter (Bjorkgren et al., 2012) to drive Cre specifically in the most proximal “initial segment” of the caput epididymis (**Figure 4A-B**). Mice carrying both the TU tracer cassette and *Defb41*:Cre exhibited the expected caput-specific expression of UPRT (**Figure 4C**). Injection of 4-TU into these animals resulted in caput-specific incorporation of 4-TU as assessed by biotinylation of total RNA (**Supplemental Figure S4A**) and by SLAM-Seq analysis of purified small RNAs (**Figure 4D**). Confident that we could reliably detect small RNAs synthesized in the caput epididymis, we therefore carried out SLAM-Seq for small RNAs isolated from the testis, caput epididymis, and cauda epididymal sperm in 4-TU-injected and control animals. Globally, we found no increase in U->C mutation in microRNAs purified from the testis, and a significant 2-fold increase in U->C in caput epididymis RNAs as expected (**Figure 4E, Table S3**).

Consistent with the hypothesis that cauda sperm carry microRNAs originally synthesized in the caput epididymis, SLAM-Seq of cauda sperm microRNAs revealed significantly greater 4-TU labelling than testes (**Figure 4E-F, Supplemental Figure S4B**). Although this effect is quantitatively modest, it is significant – **Figure 4F** shows the cumulative distribution plots for U->C mutations across all microRNAs for these samples (KS p value = 0.0003). Taken together, these data conclusively demonstrate that at least a subset of small RNAs in mature sperm are originally synthesized in the caput epididymis, and further support our hypothesis that small RNAs are trafficked from epididymis to sperm via epididymosomes.

## DISCUSSION

Our data provide a detailed view of the biogenesis of the small RNA repertoire of mature mammalian sperm. We characterized dynamics of small RNAs across sperm development, finding that multiple waves of small RNAs are synthesized/loaded into sperm over the course of testicular spermatogenesis and post-testicular maturation in the epididymis. Moreover, we show that caput epididymosomes can deliver small RNAs to testicular spermatozoa in vitro, and use chemogenetic tracking to definitively identify epididymis-derived RNAs in mature sperm in intact animals. Together, our data reveal a surprising role for soma-to-germline trafficking in sculpting the sperm RNA payload.

### Small RNA dynamics throughout sperm maturation

Analysis of small RNA repertoires of various purified sperm populations confirmed prior reports of a global loss of piRNAs and gain of tRFs that occurs over the course of post-testicular maturation, and further identified a narrow anatomical window during which this transition occurs. In addition to this large-scale remodeling of the sperm small RNA payload, we also identified more subtle dynamic behaviors for smaller subsets of RNAs. Most interestingly, a large group of microRNAs were abundant in testicular sperm populations and are known to be expressed during spermatogenesis, but were then lost in caput epididymal sperm before increasing in abundance again in cauda sperm.

These RNAs included a large number of microRNAs encoded in genomic clusters, including the imprinted X-linked miR-880 cluster, the miR-17-92 “oncomiR” cluster, and many smaller clusters. Importantly, it is highly unlikely that isolation of caput sperm somehow enriches for defective sperm undergoing a checkpointing or resorption process, as we have found in separate studies that ICSI using caput sperm and cauda sperm results in identical fertilization and blastocyst progression rates (Colin Conine and OJR, manuscript in preparation).

Understanding the basis for this unusual behavior will be of great interest. The likeliest scenario is that these microRNAs are degraded, or ejected in the residual body, along with the bulk of piRNAs upon epididymal entry. They would then either be trafficked from the epididymis to caput sperm in epididymosomes, or, less likely, generated in situ via processing of precursor transcripts. One possibility is that entry into the epididymis is accompanied by a nearly global purge of small RNAs including microRNAs, with those microRNAs that appear to maintain abundance in caput epididymis (such as various let-7 family members) simply being those that are replenished earlier than those that are gained more distally. Whatever the mechanism for this process, the reason for the strong caput/cauda gradients in expression and trafficking of various small RNAs remains unclear, and will be the subject of future studies.

### Soma-to-germline RNA trafficking in the epididymis

We also sought to characterize the mechanism responsible for the massive remodeling of the sperm RNA payload that occurs in the epididymis. Multiple lines of evidence support the hypothesis that a wide variety of small RNAs are synthesized in the epididymis epithelium and subsequently trafficked to maturing sperm in small extracellular vesicles, collectively known as epididymosomes: 1) Small RNAs that are gained as sperm enter the epididymis, such as tRFs, are highly abundant in the epididymis epithelium. 2) More locally, gradients in small RNA expression between the proximal and distal epididymis are manifest in sperm transiting the relevant region (as in, eg, the ~10-fold difference in tRF-Val-CAC between caput and cauda epididymis, sperm, and epididymosomes). 3) Using the TU tracer system, we definitively show here that microRNAs first synthesized in the caput epididymis make their way to maturing sperm, thus demonstrating that soma-to-germline transfer of small RNAs occurs in intact animals. 4) The likely mechanism for this transfer is RNA trafficking in epididymosomes, as purified epididymosomes carry a very similar RNA payload to that gained by sperm during epididymal transit. 5) Epididymosomes can deliver small RNAs to sperm in vitro.

Our data thus identify a role for soma-to-germline trafficking in shaping the sperm RNA repertoire in mammalian sperm. This unites mammalian spermatogenesis with gametogenesis in a number of other species where RNAs produced in somatic support cells have been suggested or shown to be subsequently transferred into developing germ cells (Bourc’his and Voinnet, 2010; Slotkin et al., 2009). Previously-described examples of such soma-to-germline communication include: 1) communication between the somatic macronucleus and the germline micronucleus in ciliates such as *Tetrahymena* and *Oxytricha*, 2) piRNA communication from somatic follicle cells to the germline during *Drosophila* oogenesis, and 3) release of transposon repression in vegetative nuclei in *Arabidopsis* pollen, with resulting expression of transposons leading to generation of antitransposon RNAs which are transported into the sperm nuclei. Intriguingly, the process detailed here is quite distinct from most prior examples of soma-to-germline communication, as the trafficking process involves a vesicle intermediate rather than direct cytoplasmic connections (Mahajan-Miklos and Cooley, 1994; McCue et al., 2011), and the RNAs being shipped are primarily microRNAs and tRNA fragments as opposed to repeat element-derived piRNAs/24 nt RNAs. Importantly, although a multitude of cell types secrete RNAs into extracellular vesicles in mammals (Tkach and Thery, 2016), there is as yet no evidence that circulating vesicles have access to the germline, which is protected by the blood-testis barrier – for example, we find no evidence in the *Alb*:Cre X TU-tracer experiments for liver to germline communication. Thus the epididymis, along with somatic cells of the testis such as Sertoli cells, may occupy a privileged position in this regard.

### Epigenetic remodeling during gametogenesis

Many potentially heritable “epigenetic marks” in the germline are known to be erased during key stages of development, as for example cytosine methylation patterns undergo two major waves of erasure and re-establishment during the mammalian life cycle (Feng et al., 2010) – once in primordial germ cells, and again shortly after fertilization. Our findings reveal novel epigenetic reprogramming events that occur during mammalian spermatogenesis, as we document a major transition from a piRNA-dominated stage of testicular spermatogenesis to the ultimate tRF-dominated payload of epididymal sperm, as well as a transient loss of clustered microRNAs that occurs in the caput epididymis.

The mechanism by which this epigenetic remodelling occurs is of great interest given the mounting evidence that environmental signaling may be able to modulate specific epigenetic marks in germ cells and thereby potentially affect the phenotype in future generations (Lane et al., 2014; Rando, 2012). While it is well appreciated that our dietary conditions can influence the epigenome in somatic cells (Sharma and Rando, 2017), it has not been clear how this information is communicated to the developing gametes and thus to offspring. The origin of sperm small RNAs in the epididymis provides a focus for future studies on the mechanism by which environmental conditions might influence the sperm epigenome. Understanding the molecular basis underlying sorting and packaging of small RNAs into epididymosomes will be of great interest (Gutierrez-Vazquez et al., 2013; Koppers-Lalic et al., 2014) as it will not only uncover basic mechanism of communication between the somatic cells of epididymis and the maturing gametes, but will also shed light on how environmental information is signaled to sperm.

## ACKNOWLEDGEMENTS

We thank C. Conine and other members of the Rando lab for critical reading of the manuscript and helpful comments. We thank P. Sipilä for the generous gift of the *Defb41*:Cre mouse strain. This work was supported by NIH grants R01HD080224 and DP1ES025458 (OJR), and the Austrian Academy of Sciences and the European Research Council ERC-StG-338252 (SLA). US is a Charles H. Hood postdoctoral Fellow.

## MATERIALS AND METHODS

### Mouse strains and husbandry

Wild type FVB/NJ mice obtained from Jackson Laboratory were used for all experiments other than TU-tracer experiments. For TU-tracer experiments, B6(D2)-Tg(CAG-GFP,-Uprt)985Cdoe (CAGFPstopUPRT mice) and *Albumin*-cre mice were purchased from Jackson Laboratory. *Defb41* iCre KI (TUKO 10) mouse embryos were a gift from Dr. Petra Sipilä, Univeristy of Turku Finland. *Defb41* iCre KI (TUKO 10) mouse line was regenerated at the Mouse Facility of UMass Medical School. All animal care and use procedures were in accordance with guidelines of the Institutional Animal Care and Use Committee.

### Tissues and sperm cells isolation

Testes, epididymis, caput sperm, cauda sperm, spermatocytes and round spermatids were isolated as described previously (Sharma et al., 2016). For mature testicular spermatozoa isolation, testes from one mouse (8-12 weeks old) were minced in a 35-mm Petri dish containing 1ml 0.9% NaCl. Finely minced tissue slurry was then transferred to a 15ml conical tube and set aside for 3-5 minutes to allow tissue pieces to settle down. Next, the cell suspension was loaded onto 10.5 ml of 52% isotonic percoll (Sigma). The tubes were then centrifuged at 12000X g for 10 minutes at 10^°^C. The pellet was resuspended in 10 ml of 0.9% NaCl and spun at 600X g at 4^°^C for 10 minutes. The pellet was next washed with 0.45% NaCl solution 3 times, followed by one wash with 1X PBS to retrieve purified mature testicular spermatozoa. Spermatozoa pellets were then either flash frozen or used for fusion with epididymosomes.

### Epididymosome isolation

Epididymosomes were isolated as previously described (Sharma et al., 2016). Briefly, caput epididymides were dissected from 8-12 weeks old male mice and placed in dishes containing 1 ml Whitten’s media (100 mM NaCl, 4.7 mM KCl, 1.2 mM KH2PO4, 1.2 mM MgSO4, 5.5 mM Glucose, 1 mM Pyruvic acid, 4.8 mM Lactic acid (hemicalcium), and HEPES 20 mM) pre-warmed at 37°C. The epididymides were then gently squeezed using forceps to isolate epididymal luminal content. The dishes containing epididymal luminal contents were then placed in an incubator set at 37°C with 5% CO2 for 15 minutes, to allow any remaining epididymal contents to release from the tissue. Next, the media containing epididymal luminal contents was transferred to a 1.5 ml tube and allowed to incubate for an additional 15 minutes. At the end of the 15 minutes, any tissue pieces or non-motile sperm settled down at the bottom of the tube and all the contents of the tube except for the bottom ~50 μl were transferred to a fresh tube. Next, the tube was spun in a tabletop centrifuge at 2000X g for 2 minutes to pellet down sperm. Supernatant, which contains epididymosomes, was then transferred to a fresh tube and centrifuged at 10000X g for 30 minutes at 4°C to get rid of any non-sperm cells and cellular debris. Supernatant from this spin was then transferred to a polycarbonate thick wall tube (13 X 56 mm, Beckman Coulter, Catalog number 362305) and centrifuged at 100000X g for 2 hours at 4°C in a tabletop ultracentrifuge (Beckman Optima TL) using TLA100.4 rotor. The pellet from this spin was then washed with 500 μl 1X PBS and centrifuged for another 2 hours at 100000X g at 4°C. Finally, the pellet containing epididymosomes was resuspended in 50 μl ice-cold 1X PBS, transferred to a 1.5 ml tube and flash frozen in liquid nitrogen or used for fusion with sperm.

### Testicular spermatozoa and caput epididymosomes fusion

After final wash with 1XPBS, caput epididymosomes were resuspended in 62.5 μl media out of which 12.5 μl was flash frozen and 50 μl was used for fusion with testicular spermatozoa. For fusion reaction, purified 100 μl of testicular spermatozoa were mixed with 50 μl of caput epididymosomes in presence of 1 mM ZnCl2 and incubated at 37°C for 2 hours. In parallel, a mock fusion reaction was also performed where no epididymosomes were added to the reaction. At the end of the incubation, additional 850 μl of 1X PBS was added to the reaction and sperm were spun down at 4000X g for 5 minutes. Next, the pellet was washed with 1X PBS three times and processed further for RNA extraction.

The finding of epididymosomal RNAs in resulting “reconstituted sperm” (**Figure3**) show that caput epididymosomes are either capable of fusing with mature testicular spermatozoa to deliver their small RNA cargo, or, less likely, that a small subset of vesicles with a distinctive small RNA payload adhere to spermatozoa strongly enough to resist removal by several consecutive washing steps

### TaqMan assays

Taqman assays were performed as described previously (Sharma et al., 2016). Custom small RNA assays for tRF-Val-CAC, tRF-GluCTC, tRF-GlyGCC and pre-designed assay for miR34c were purchased from Life Technologies.

### Small RNA deep sequencing

Isolation of 18-40 nts small RNAs was carried out as previously described (Sharma et al., 2016) by purification from 15% polyacrylamide-7M urea denaturing gels. The resulting small RNA was used for preparing sequencing libraries using the Small RNA TrueSeq kit from Illumina (as per the manual). Small RNA sequencing data was analyzed as described previously (Sharma et al., 2016).

### Generation of mice expressing tissue-specific UPRT

To generate mice expressing tissue specific UPRT, we crossed homozygous CAGFPstopUPRT mice with either liver-specific *Albumin*-cre mice or caput epididymis specific *Defb41*-cre mice, to generate heterozygous [*Uprt-/+, Alb-Cre*] or [*Uprt-/+, Defb41-Cre]* mice, respectively. Since 4-thiouracil incorporation is sensitive to the levels of UPRT, we then backcrossed these mice with homozygous CAGFPstopUPRT mice to produce mice homozygous for *Uprt* gene. Mouse genotyping was performed using quantitative PCR (qPCR) with DNA isolated from ear punches. For genotyping CAGFPstopUPRT mice, following primers and probes were used: 5’-ATT CCA AGA TCT GTG GCG TC-3’ and 5’-CTT CTC GTA GAT CAG CTT AGG C-3’ for mutant allele; 5’-CAC GTG GGC TCC AGC ATT-3’ and 5’-TCA CCA GTC ATT TCT GCC TTT G-3’ for wildtype allele; probe 5’-/Cy5/CCA ATG GTC GGG CAC TGC TCA A/Black Hole Quencher 2/-3’ for wildtype allele and probe 5’-/6-FAM/CCG CAT CGG GAA AAT CCT CAT CCA/Black Hole Quencher 2/-3’ for mutant allele. *Albumin*–cre mice were genotyped using: 5’-TTG GCC CCT TAC CAT AAC TG-3’ for both wildtype and mutant allele, 5’-TGC AAA CAT CAC ATG CAC AC-3’ for wildtype allele, and 5’-GAA GCA GAA GCT TAG GAA GAT GG-3’ for mutant allele. Primers for *Defb41*-cre mice included: 5’-AGA TGC CAG GAC ATC AGG AAC CTG-3’ and 5’-ATC AGC CAC ACC AGA CAC AGA GAT C-3’ for transgene, 5’-CTA GGC CAC AGA ATT GAA AGA TCT-3’ and 5’-GTA GGT GGA AAT TCT AGC ATC ATC C-3’ for internal positive control. The qPCR conditions were as follows: 95°C for 3 min, followed by 10 cycles of 95°C for 30 sec, 65°C for15 sec, decrement of temperature by 0.5°C per cycle, 68°C for10 sec, and 28 cycles of 95°C for 20 sec, 60°C for 20 sec, and 72°C for 15 sec, with a final extension at 72°C for 2 min.

### 4-thiouracil (4-TU) treatment

4-TU was dissolved in DMSO at 200 mg/ml and stored at -20°C. Prior to use, the stock solution was diluted at 1:4 with corn oil with rigorous shaking and working solution was then kept in dark. 8-12 weeks old adult mice were intraperitoneally injected with either 4-TU (400 mg/kg body weight) or solvent alone (control) every other day for 5 times, as sperm transits from caput to cauda epididymis in approximately 10 days. Tissue and sperm samples were collected (as described above) 5 hours after the last injection, flash frozen in liquid nitrogen and stored at -80°C until RNA extraction.

### Northern and Dot blot analysis

We followed published method for dot blot analysis with some modifications (Radle et al., 2013). In brief, equal amount of RNA (in binding buffer) from different experimental groups, was applied to Zeta membrane (Bio-Rad). A biotin-labeled DNA oligo was used as a positive control. The membrane was air-dried at room temperature for 5 minutes, followed by blocking at room temperature for 30 minutes with 10% SDS in PBS and 1mM EDTA. Next, the membrane was incubated in a solution containing 10% SDS in PBS and Streptavidin-HRP (Thermo Fisher Scientific) for 15 minutes at room temperature, followed by three washes with PBS containing SDS for 5 minutes each. After the final wash, biotinylated RNA were detected with Amersham ECL Western Blotting Reagent (GE Healthcare Life Sciences) using Amersham Imager 600 (GE Healthcare Life Sciences). For Northern blot analysis, 30 μg of RNA was denatured and run on a 15% acrylamide 7M urea gel, followed by transfer to a positively charged nylon membrane and UV-crosslinking. The membrane was next incubated with Streptavidin-HRP, washed and detected using ECL reagent, as above.

### Western blot

Protein extracts were prepared from mouse tissues using RIPA lysis buffer (150 mM sodium chloride, 1.0% NP-40, 0.5% sodium deoxycholate, 0.1% SDS, and 50mM Tris, pH 8.0). Protein extracts were denatured with 2X Laemmli buffer (4% SDS, 10% 2-mercaptothanol, 20% glycerol, 0.004% bromophenol blue, 0.125 M Tris-HCl, pH 6.8) at 95°C for 5 min and run on a 10% SDS-PAGE, followed by transfer to PVDF membrane. The membrane was next probed with anti-GFP (Invitrogen), anti-HA (Thermo Scientific) or anti-Actin (Thermo Scientific) antibodies at 1:1000 dilution followed by secondary antibody treatment at 1:2000 dilution (anti-mouse HRP and anti-rabbit HRP, Cell Signalling).

### SLAM-seq

4 μg of isolated total RNA was treated with 10 mM iodoacetamide under optimal reaction conditions (50 mM NaPi, pH8; 50% v/v DMSO; 15 min at 50°C). The reaction was quenched by addition of 10 mM DTT, followed by ethanol precipitation and small RNA library preparation as described previously (Jayaprakash et al., 2011). Briefly, total RNA was size selected (18-30 nt) by 15% denaturing polyacrylamide gel electrophoresis and subjected sequentially to 3’ and 5’ adapter ligation (employing linker oligonucleotides with four random nucleotides at the ligation interface to avoid ligation biases). Gel-purified ligation products were reverse transcribed using SuperScript III (Invitrogen) according to the manufacturer’s instructions, followed by PCR amplification and high-throughput sequencing on an Illumina HiSeq 2000 instrument (SR100 mode).

### Data analysis

For the analysis of SLAM-seq datasets, small RNA reads were aligned using bowtie v.1.2 (Langmead et al., 2009), allowing for 3 mismatches, to all mouse pre-miRNA hairpins described in mirbase v.21 (Kozomara and Griffiths-Jones, 2014) and extended downstream by 20 basepairs. All reads mapping to 5p and 3p arms of a miRNA hairpin locus were filtered to only retain the most frequent 5’ isoform. Mutation rates were determined by investigating the occurrence of any given mutation within the miR body (nt 1-18) of the most abundant miRNA 5´ isoforms, normalized to the genome matching instances at this position and frequency of each nucleotide within the inspected region. Reads containing T>C conversions within the miR body were counted and normalized to U-content and total mapping reads to assess abundance in cpm.

## SUPPLEMENTAL MATERIALS

**Supplemental Figures S1-S4**

**Supplemental Tables S1-S3**

## SUPPLEMENTAL TABLES

**Supplemental Table S1: Small RNA-Seq across sperm development**. All small RNA-Seq data for primary spermatocytes, round spermatids, testicular spermatozoa, and caput and cauda epididymal sperm, as indicated, as well as cauda epididymosomes. Total read numbers are provided here – for analysis throughout the manuscript, rRNA-mapping reads were discarded and remaining reads were normalized to ppm.

**Supplemental Table S2: Small RNA-Seq for epididymosomal reconstitution experiments**. As in **Table S1**, for testicular sperm either mock-treated or fused with caput epididymosomes. Note that 2 columns here (mock-treated sperm) are reproduced from **Table S1**.

**Supplemental Table S3: SLAM-Seq dataset**. Sheets show analysis of each mutation type (C>A, etc.). In each sheet, data for individual microRNAs are shown as rows, with six columns showing raw read counts for the six libraries (testis, caput epididymis, and cauda sperm, all with or without 4-TU injection), and the following six columns showing the average mutation rate for all relevant nucleotides (eg all cytosines for the C>A sheet) in the microRNA’s first 18 nt. For the T>C sheet, columns are also included with data normalized to parts per million, and three columns are included showing the 4-TU/no TU ratio for the relevant mutation rates.

## SUPPLEMENTAL FIGURE LEGENDS

**Supplemental Figure S1:**
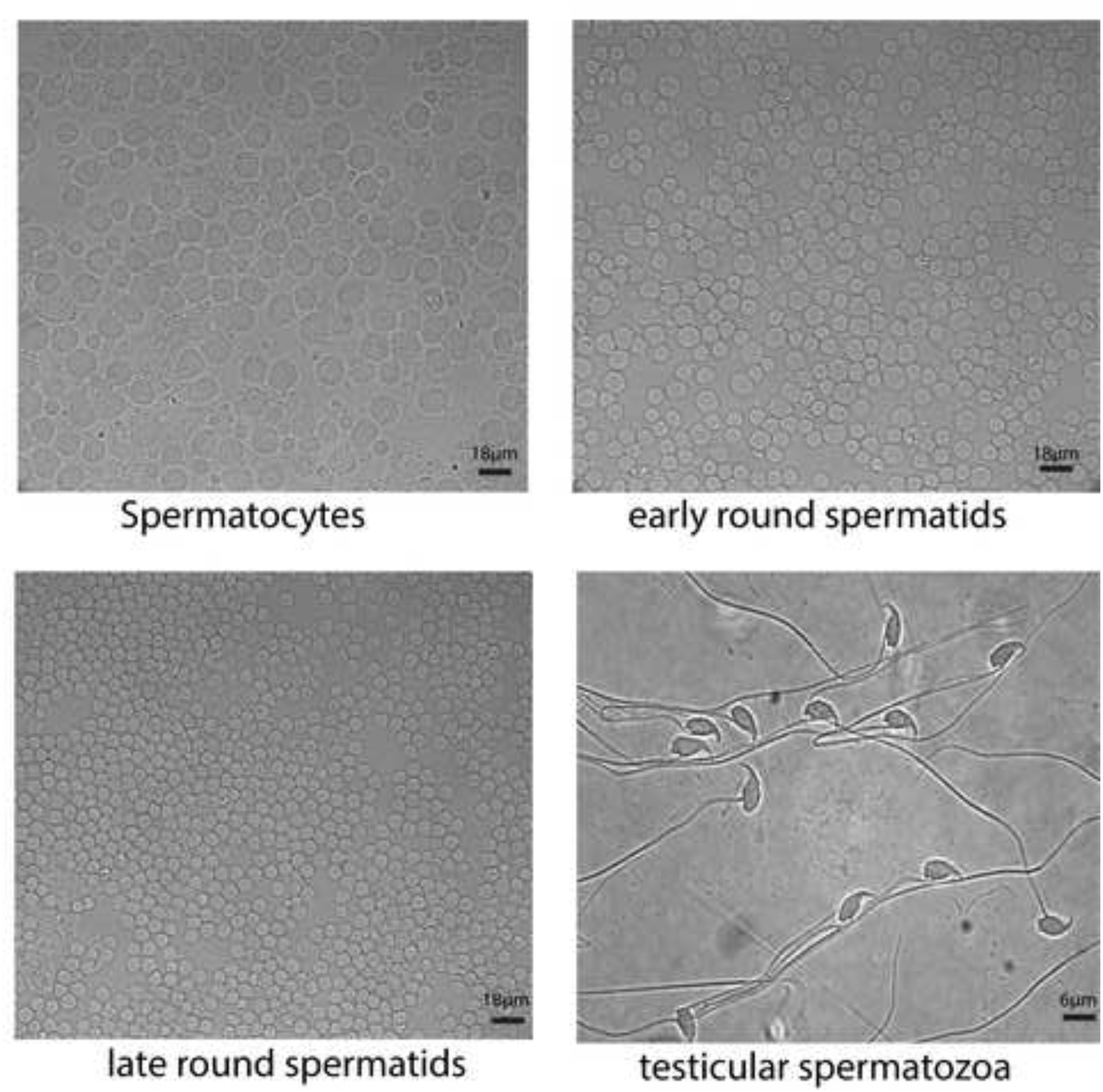
Purification of testicular spermatozoa. DIC images of purified testicular germ cells: spermatocytes, early and late round spermatids (scale bar= 18 μm), and mature testicular spermatozoa (scale bar = 6 μm).

**Supplemental Figure S2:**
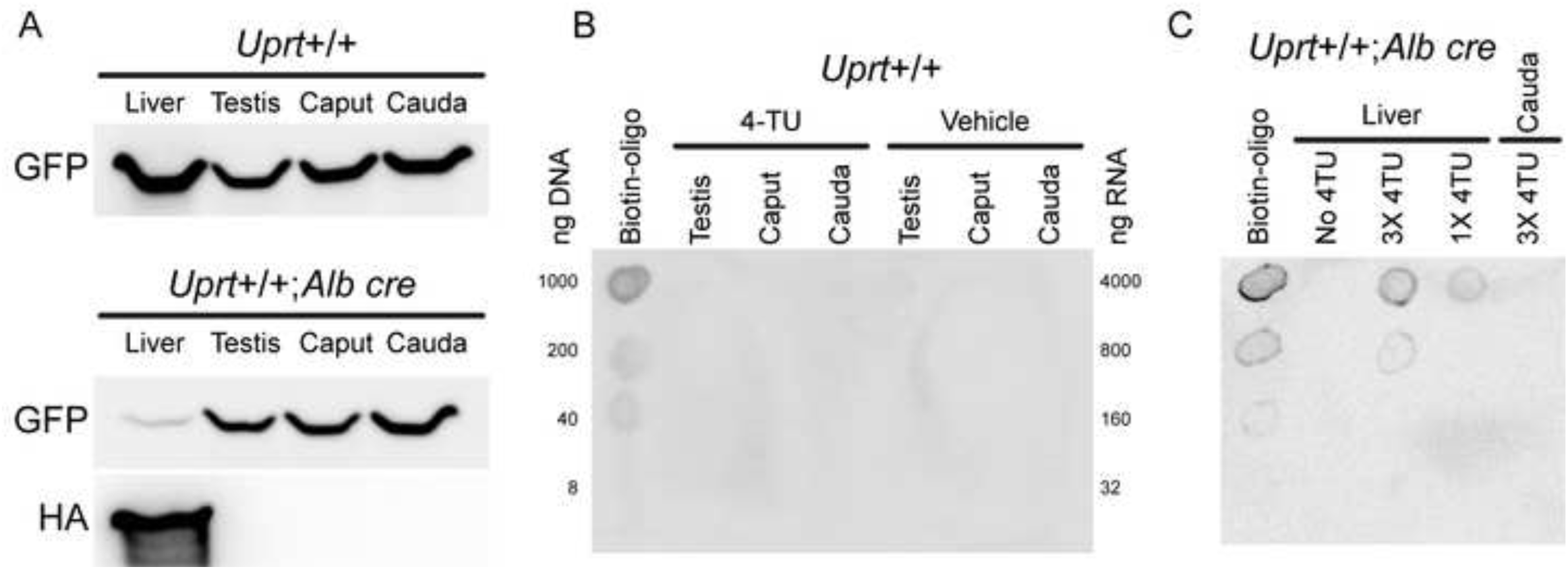
Validation of TU-tracer system. A) Western blot showing GFP and HA-UPRT levels in livers and several reproductive tissues isolated from TU-tracer animals not expressing Cre (top) or expressing liver-specific *Albumin*-Cre recombinase (bottom). In the absence of Cre, GFP is expressed from a ubiquitous promoter, with no UPRT expression. In mice expressing liver specific Cre recombinase, GFP is eliminated by LoxP recombination, leading to expression of HA-UPRT specifically in liver tissues. B) Dot blot for RNA isolated from TU-tracer animals not expressing Cre and injected with either 4-TU or vehicle alone. As shown, only positive control (Biotinylated DNA oligo) is detectable upon incubation with Streptavidin-HRP. C) Dot blot for RNA isolated from *Albumin-Cre* X TU tracer mice injected with two different doses of 4-TU. Upon 4-TU injection RNAs are specifically labeled, in a dose dependent manner, in liver tissue and not in control tissue (cauda epididymis).

**Supplemental Figure S3:**
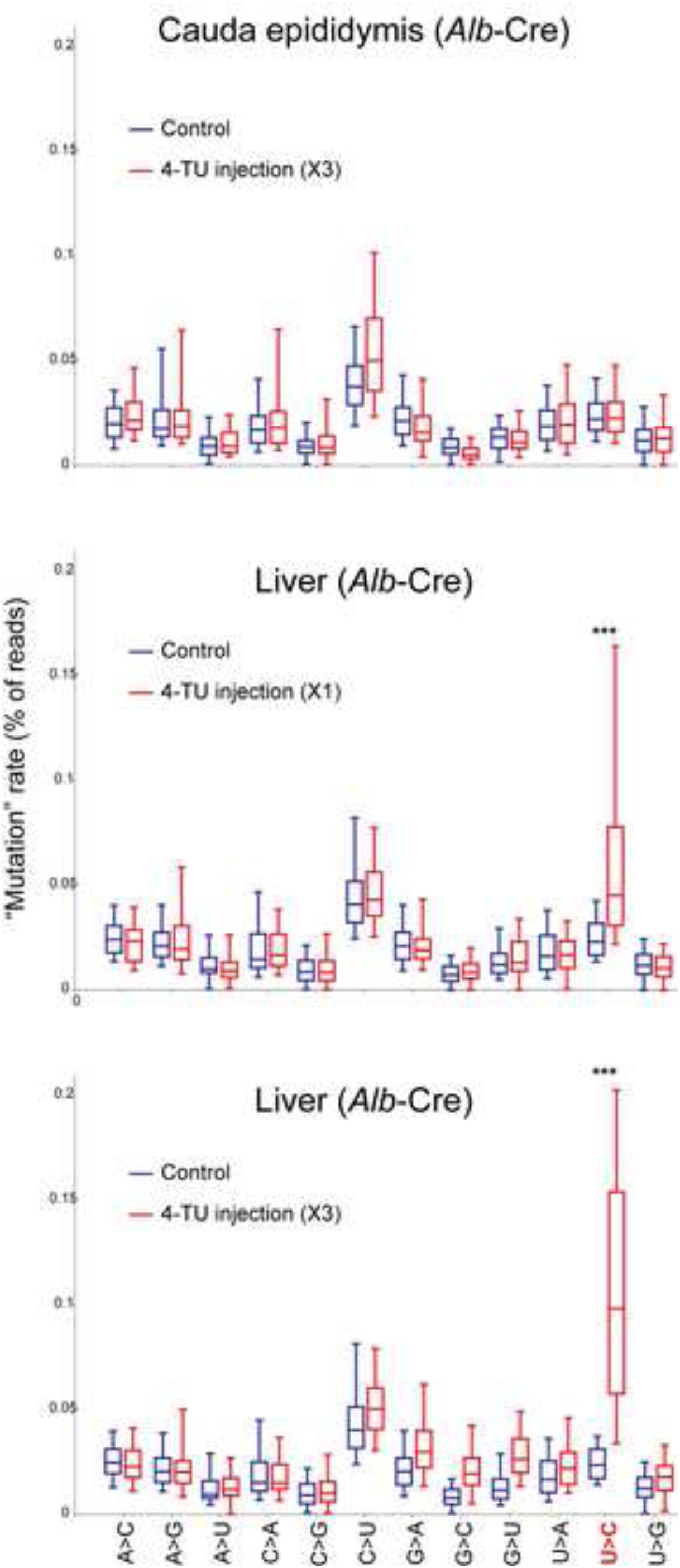
SLAM-Seq mutation rates. TU tracer X *Alb*:Cre animals were either injected with 4-TU (400 mg/kg body weight) or with vehicle (solvent only) and sacrificed 5-6 hours after the last injection for tissue harvest. Small RNAs were isolated from either cauda epididymis or from liver, and subject to SLAM-Seq, and sequencing reads were mapped to microRNAs using an error-tolerant pipeline. Mismatches were identified for all reads mapping to a given microRNA, and boxplots (box: 25^th^/50^th^/75^th^ percentile; whiskers: 5^t^^h^/95^th^ percentile) show the frequency of various mismatches for mock-injected and 4-TU-injected animals, as indicated.

**Supplemental Figure S4:**
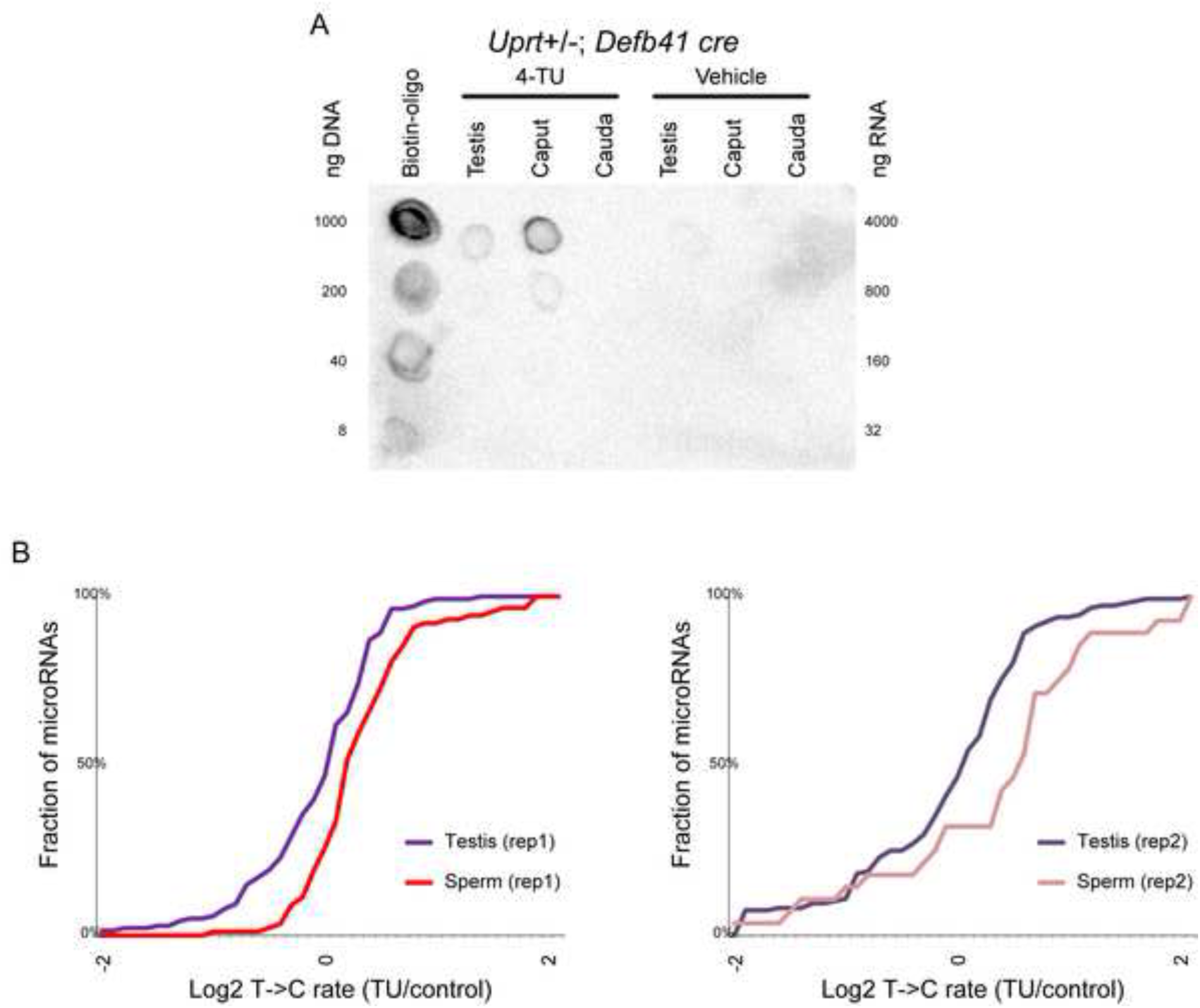
A) Dot blots for RNA isolated from TU-tracer animals expressing *Defb41-Cre* mice, either injected with 4-TU or vehicle alone. Only upon 4-TU injection, RNAs are specifically labeled in caput epididymis and not in control tissues (testis and cauda epididymis) B) Left panel reproduces the cumulative distribution plot from **Figure 4F**, right panel shows the same plot for a second replicate dataset (KS p value = 0.0027).

